## Supplementary Figures and Tables for "Tig1 regulates proximo-distal identity during salamander limb regeneration"

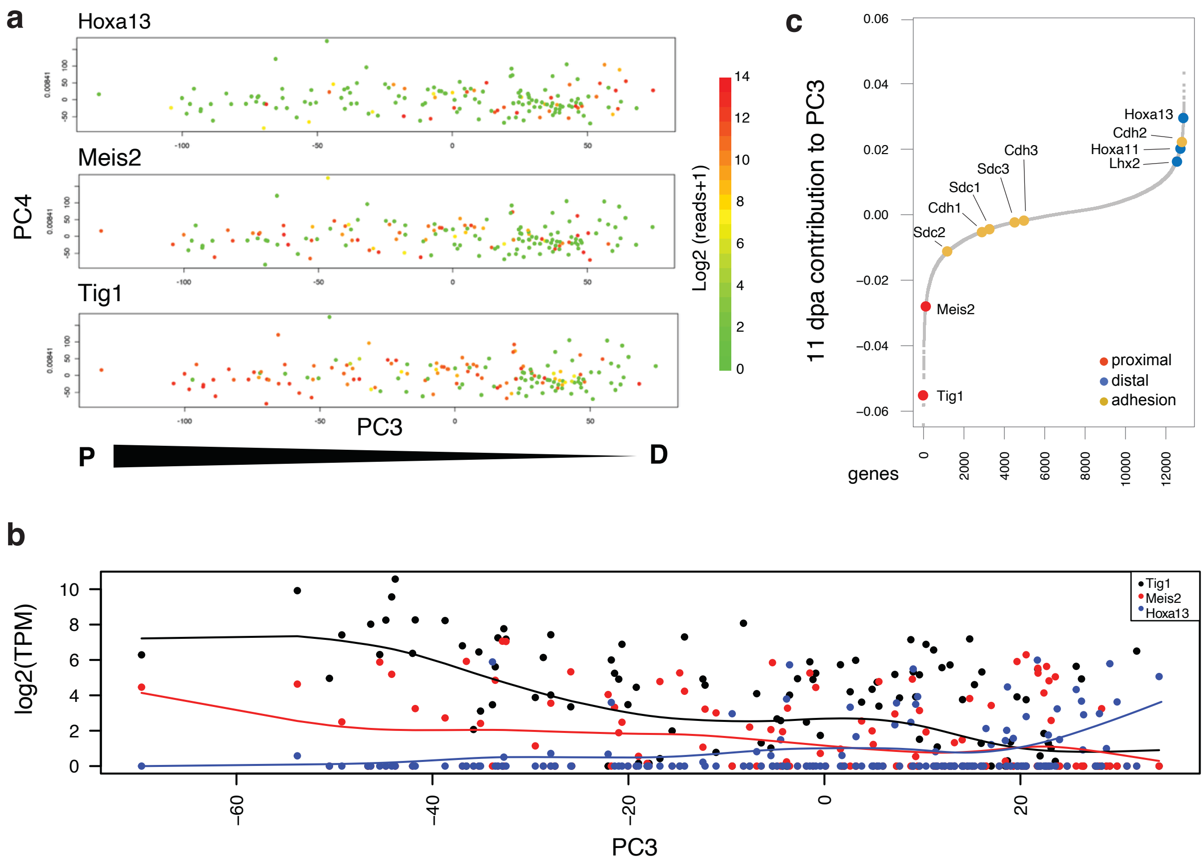


Supplementary Fig. 1

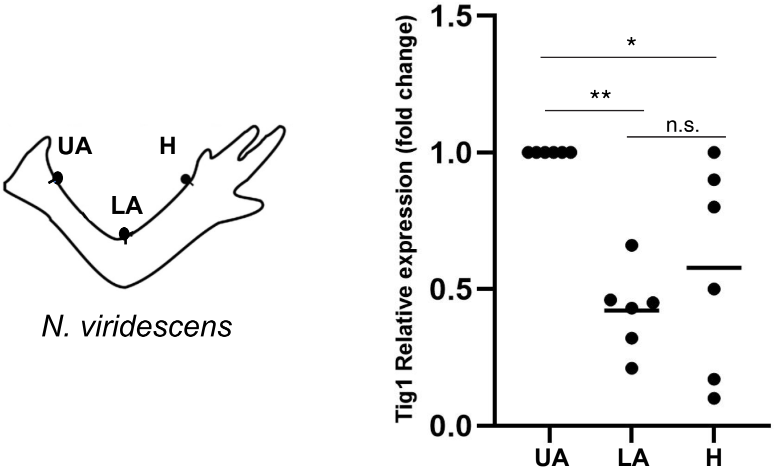


Supplementary Fig. 2

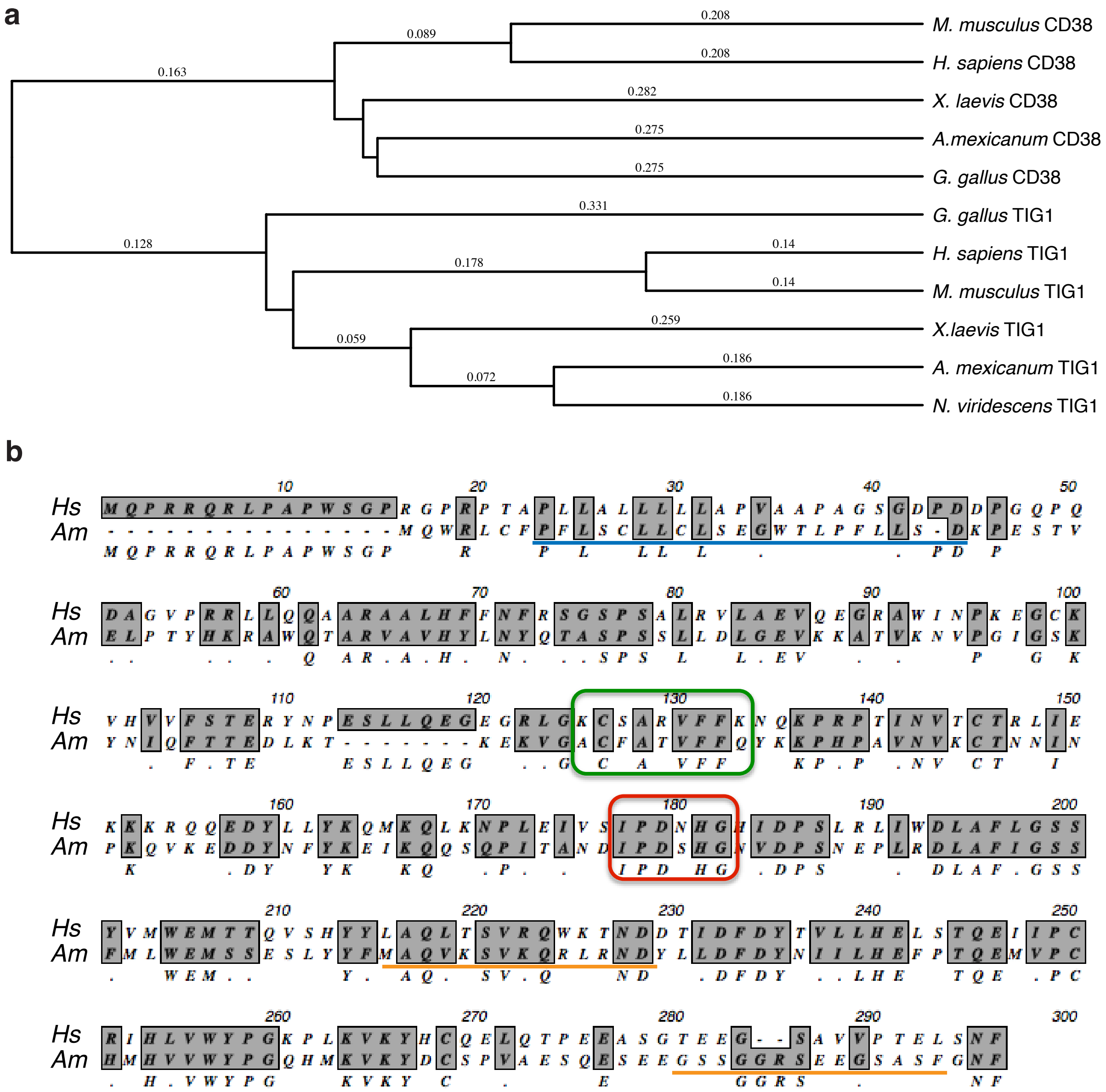


Supplementary Fig. 3

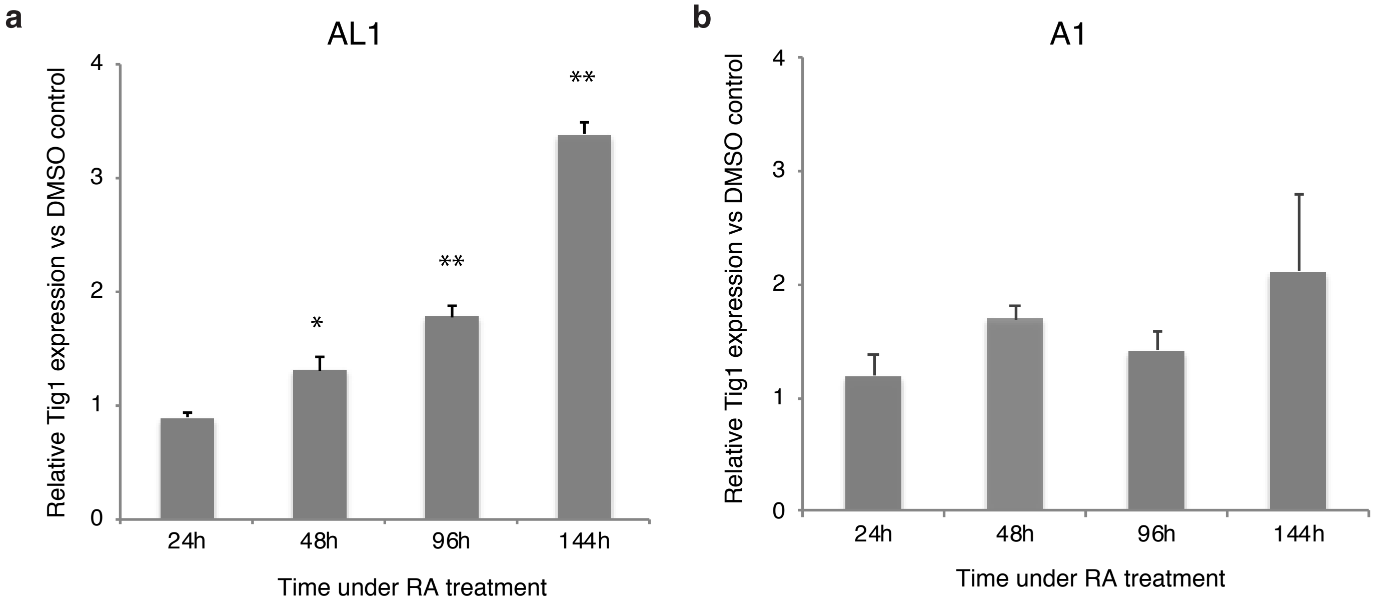


Supplementary Fig. 4

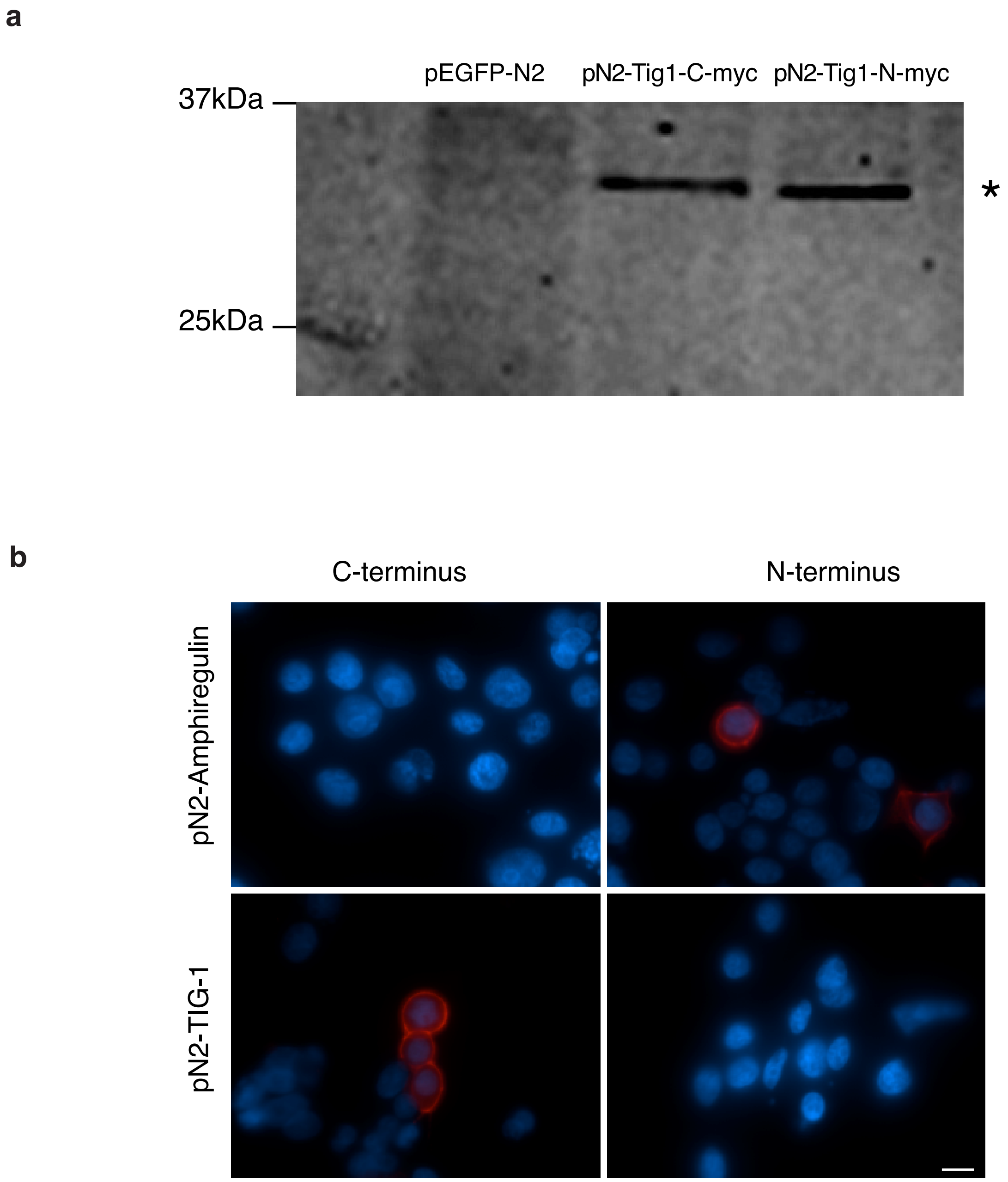


Supplementary Fig. 5

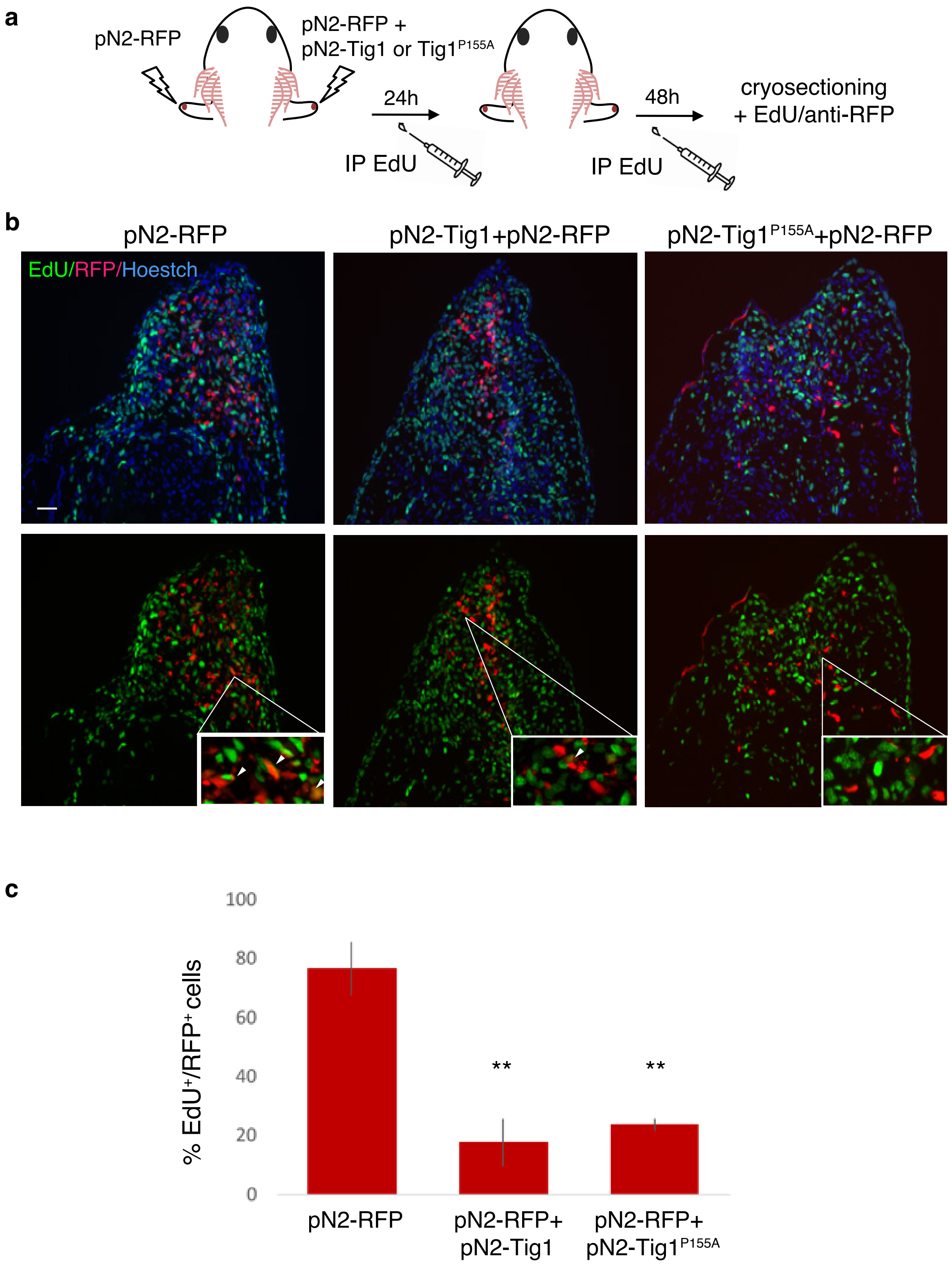


Supplementary Fig. 6

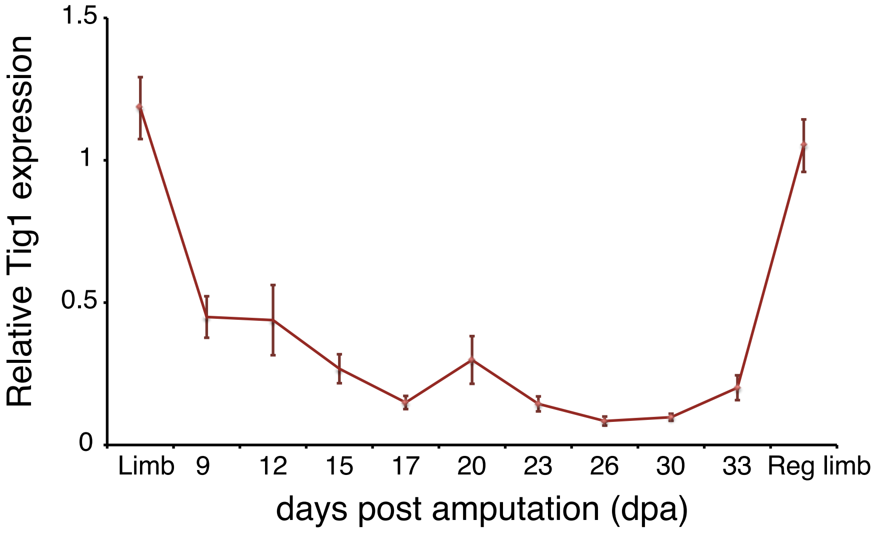


Supplementary Fig. 7

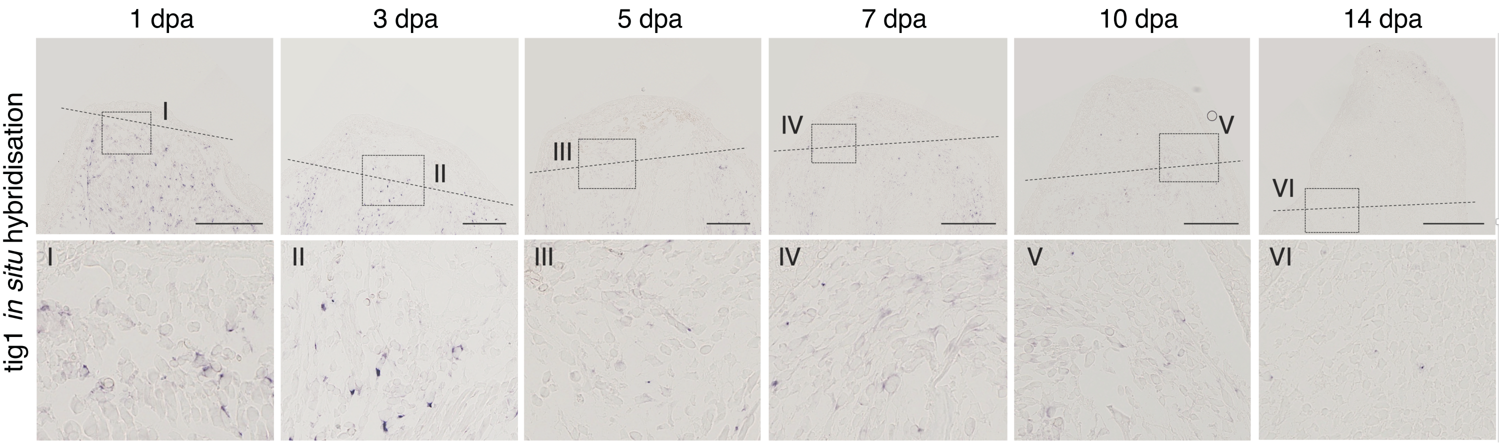


Supplementary Fig. 8

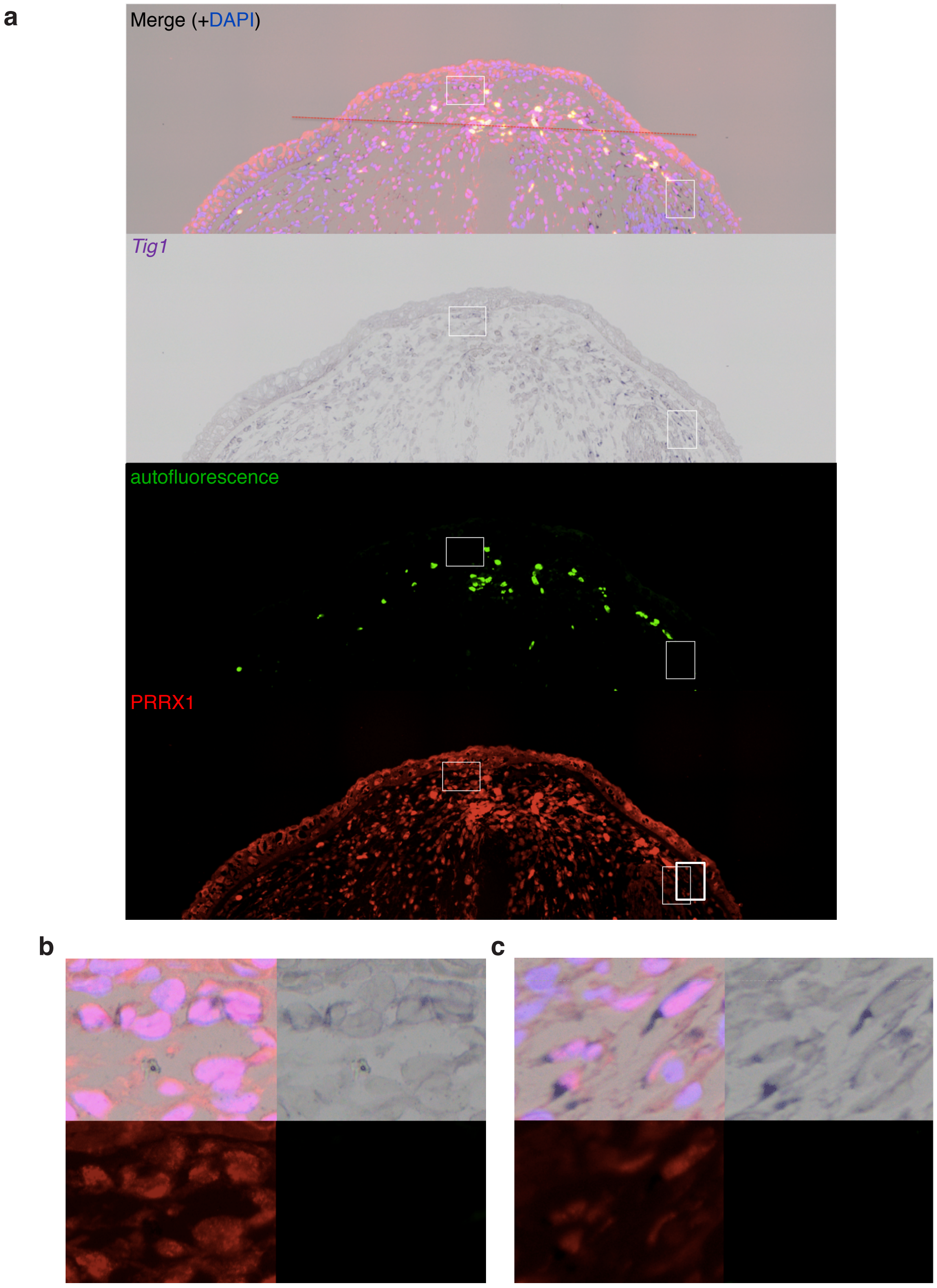


Supplementary Fig. 9

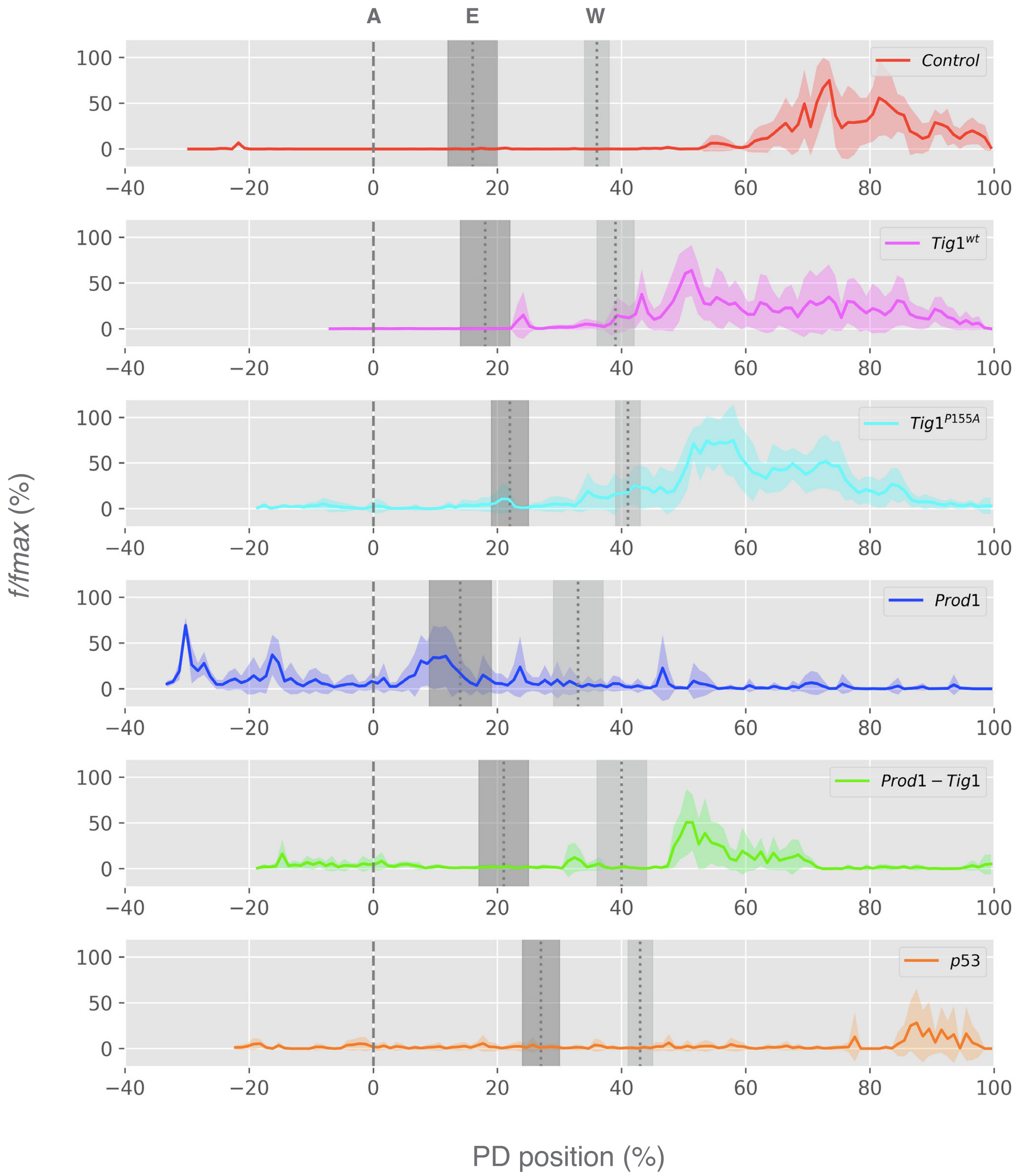


Supplementary Fig. 10

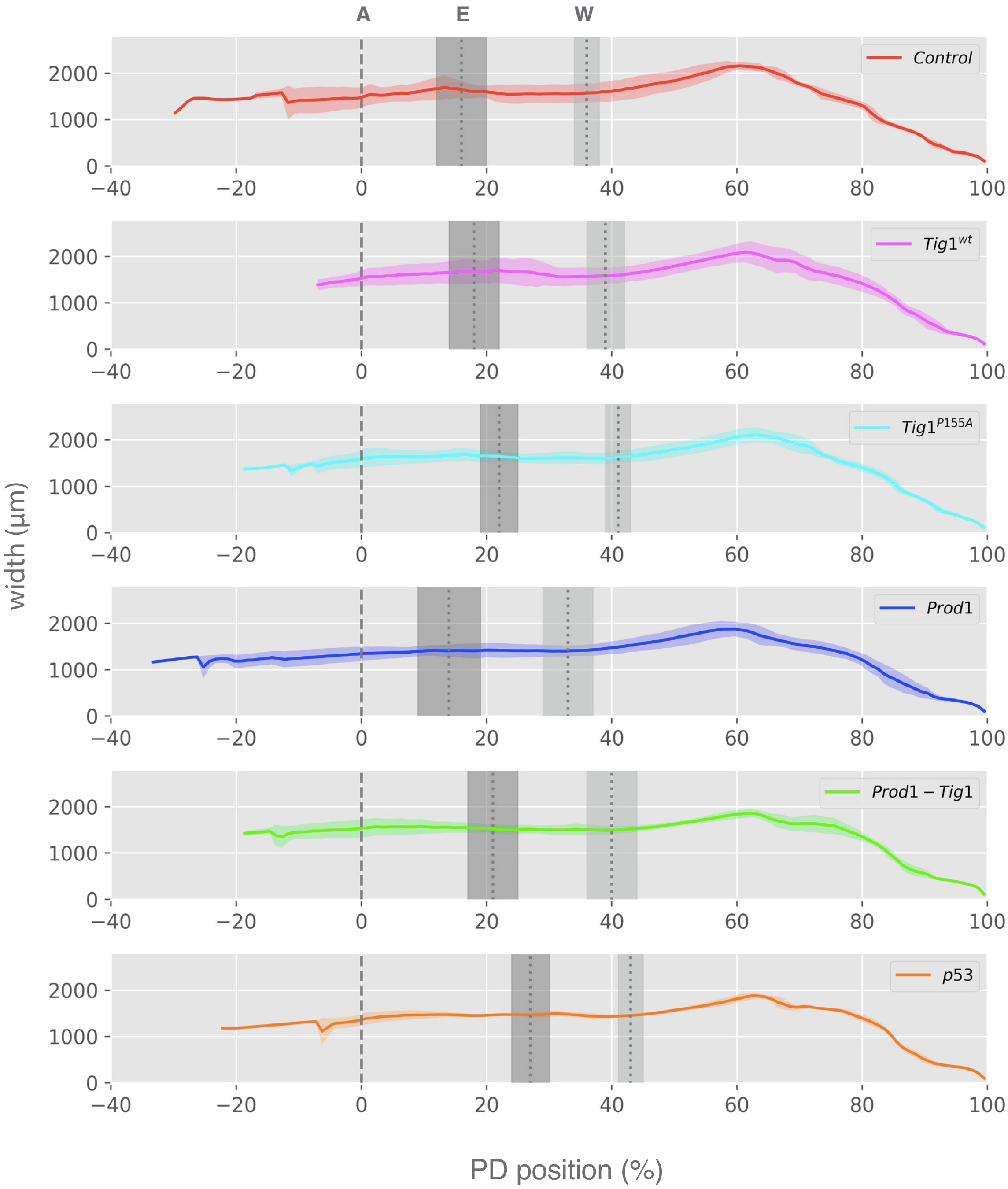


Supplementary Fig. 11

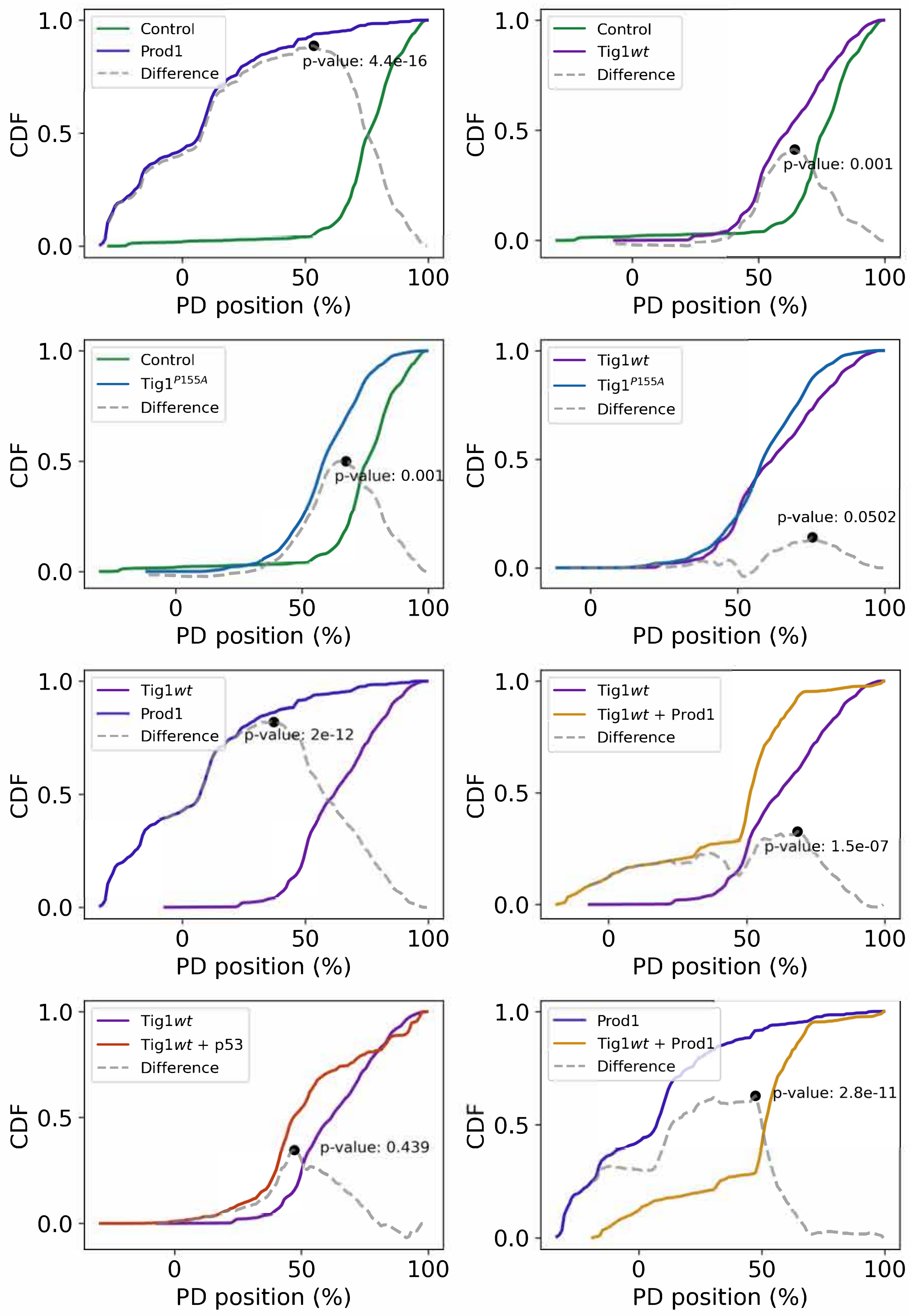


Supplementary Fig. 12

**a**

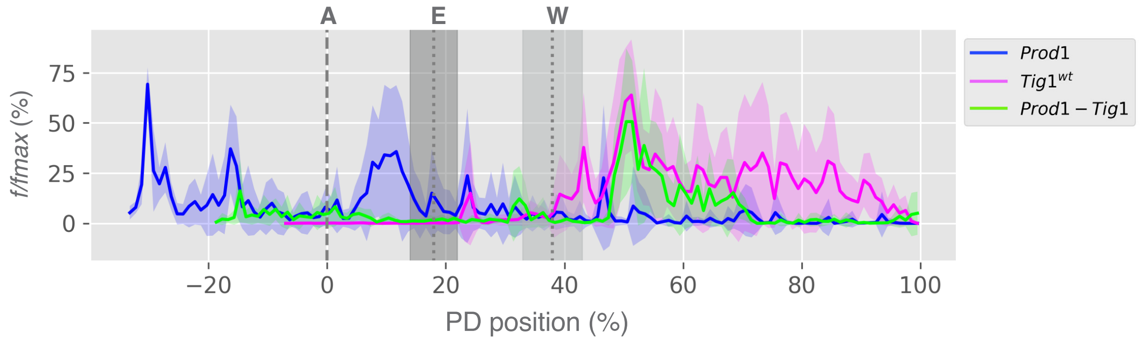


**b**

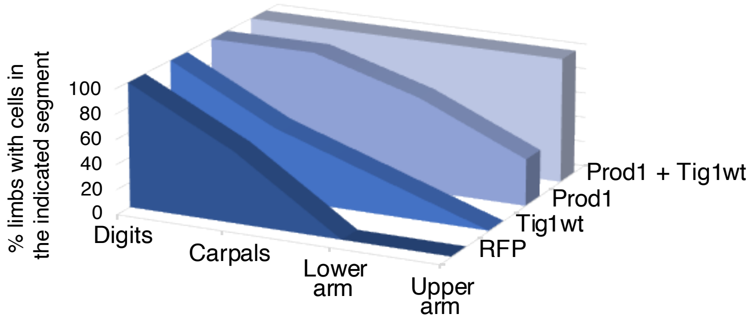


Supplementary Fig. 13

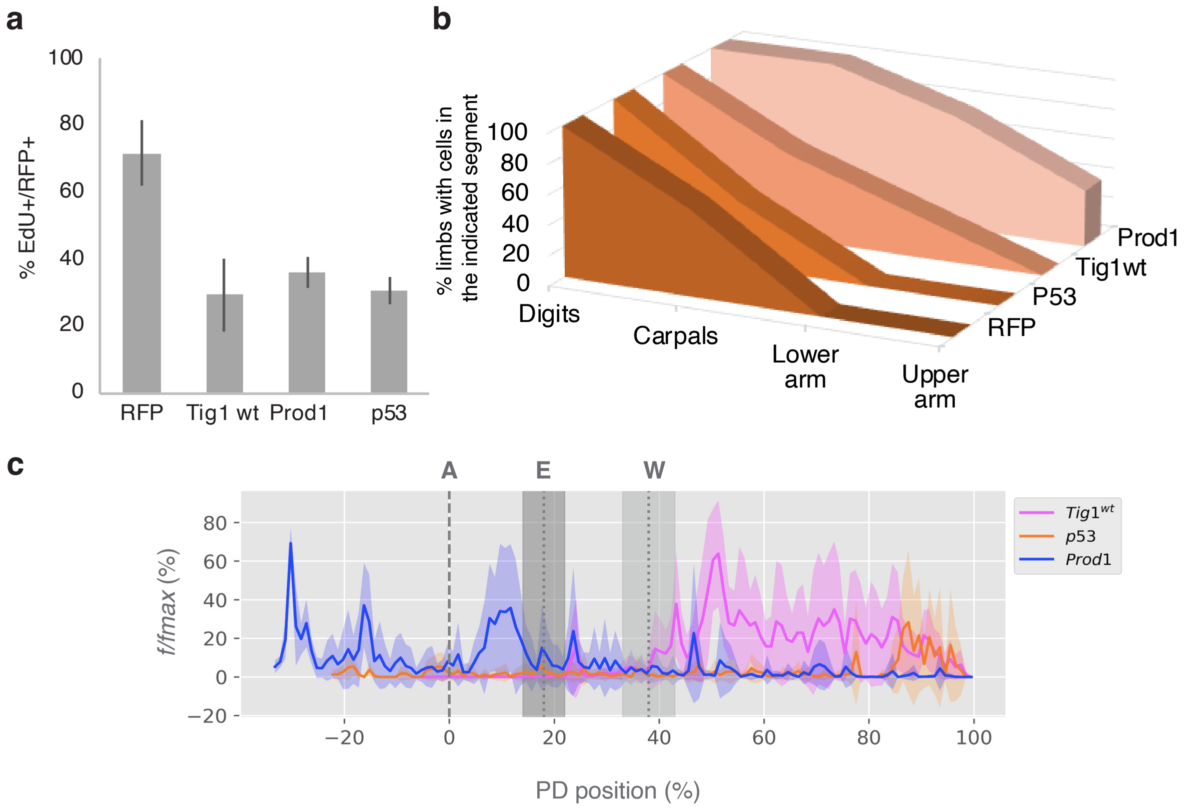


Supplementary Fig. 14

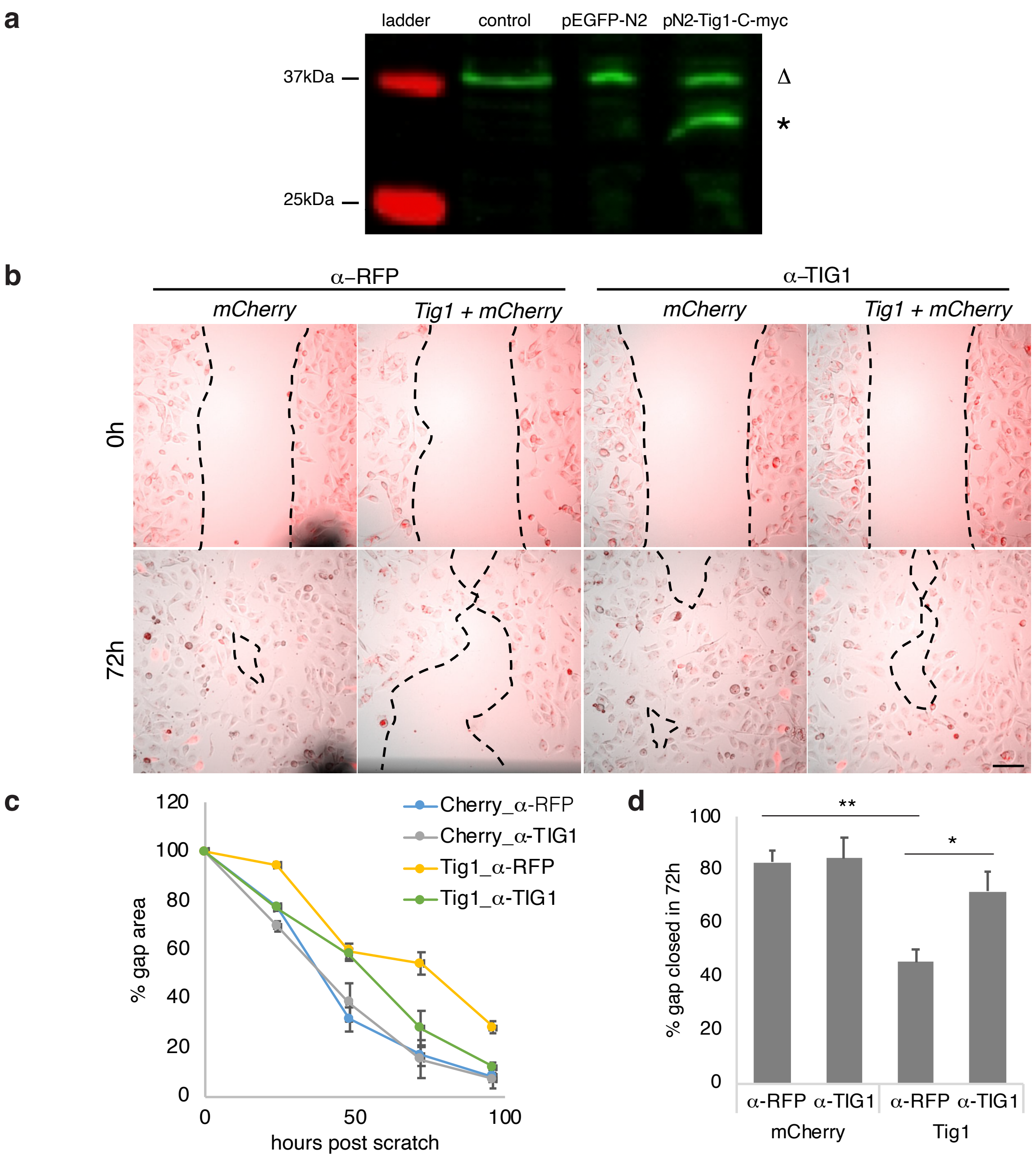


Supplementary Fig. 15

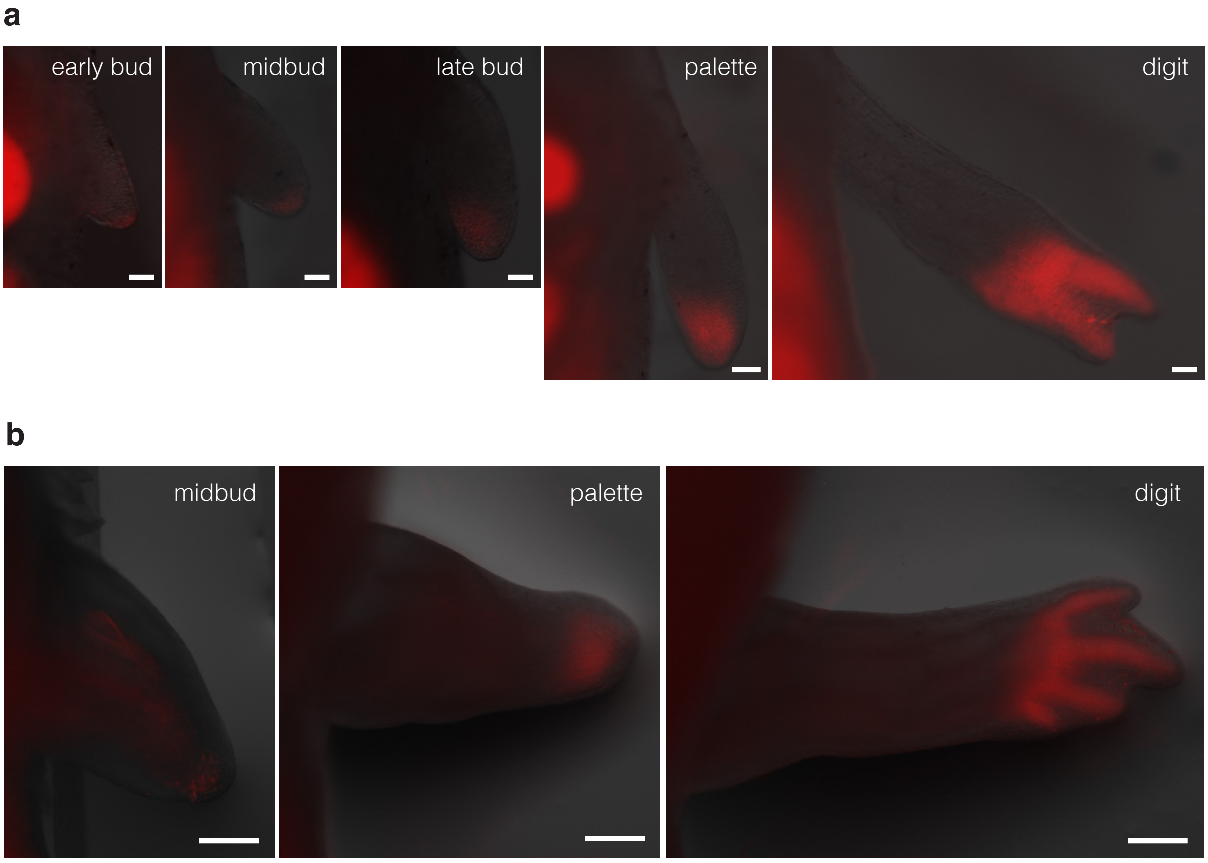


Supplementary Fig. 16

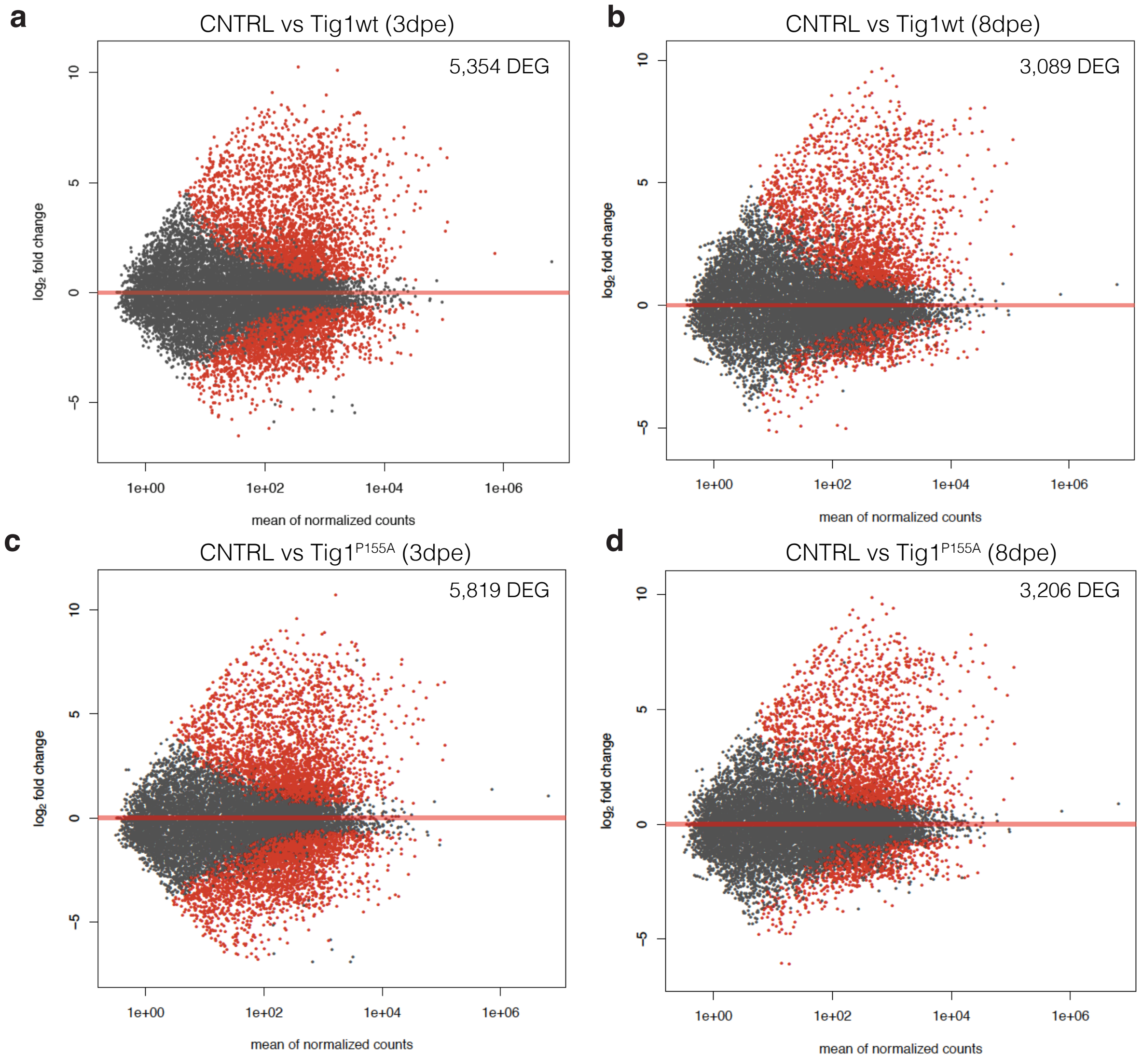


Supplementary Fig. 17

**a**

GO Analysis - Cluster B

| adj.Pval | nGenes | Pathways |
|---|---|---|
| 8.50E-90 | 325 | Immune system process |
| 1.40E-80 | 266 | Immune response |
| 6.30E-74 | 203 | Cell activation |
| 4.50E-71 | 188 | Leukocyte activation |
| 6.90E-58 | 128 | Myeloid leukocyte activation |
| 1.70E-54 | 165 | Immune effector process |
| 3.50E-51 | 189 | Regulation of immune system process |
| 2.50E-48 | 120 | Leukocyte activation involved in immune response |
| 1.40E-44 | 125 | Leukocyte mediated immunity |
| 1.50E-44 | 102 | Leukocyte degranulation |

**b**

GO Analysis - Cluster C

| adj.Pval | nGenes | Pathways |
|---|---|---|
| 5.60E-18 | 14 | Muscle filament sliding |
| 5.60E-18 | 14 | Actin-myosin filament sliding |
| 1.30E-17 | 60 | Tissue development |
| 2.60E-15 | 17 | Actin-mediated cell contraction |
| 1.90E-13 | 17 | Actin filament-based movement |
| 4.80E-13 | 13 | Myofibril assembly |
| 7.30E-12 | 31 | Actin filament-based process |
| 9.30E-12 | 40 | Epithelium development |
| 1.00E-11 | 31 | Epithelial cell differentiation |
| 1.00E-11 | 16 | Striated muscle cell development |

Supplementary Fig. 18

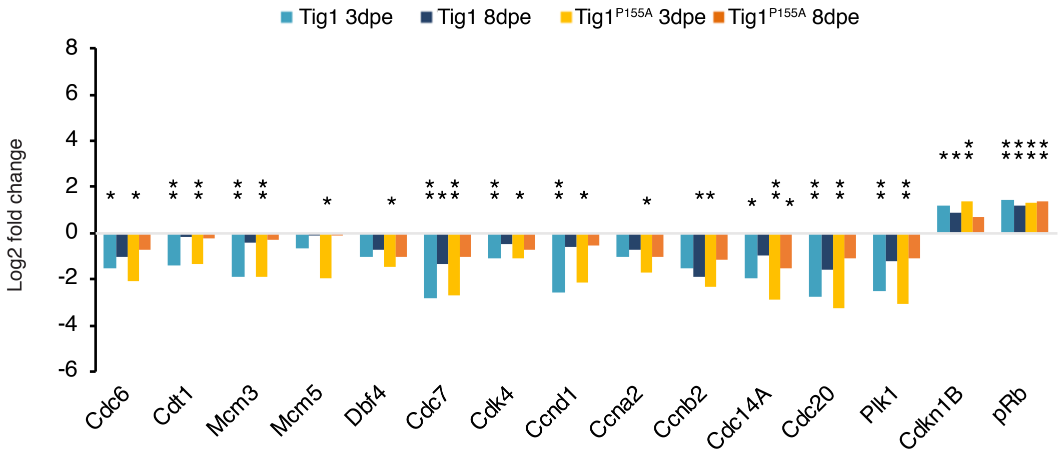


Supplementary Fig. 19

| gene | Forward (5' primer) | Reverse (3'primer) |
|---|---|---|
| Rlp4 | TGAAGAACTTGAGGGTCATGG | CTTGGCGTCTGCAGATTTTTT |
| Hoxa13 | ACTGGTTCATGAAGACAATTGCCT | CTGTTTTCACCGTCTTCCCCATTA |
| Hoxa11 | GCCAAACAGGCTTGGTTTTA | CTTCCGGCGATTGTTTAGTC |
| Hoxa9 | GAGACAAGCCTGCCATTGAC | GGTTCTGGAACCAGATCTTGAC |
| Meis1 | GCCGTCGCCAACAATGATTT | GTGATGCGTACGGTTGATGC |
| Enpp2 | CGATACCAGTCCCAACGCAT | GAGTTGCTCAATGTCCCGGA |
| Epha7 | CGTTACAATCGGCATGAGGG | GCTAGGCCTTACTTCCCGTG |
| Lhx2 | CCTCTTTCAAACACCACCAGTTG | TGTGTTTTCTTGGCGTAGGAGAT |
| Prod1 | GATCTCAAAGGTCGGTTCAGA | CACAAGCCTGCTACTTCTAGA |
| Tig1CDS | TCCAATGAGCCCCTGAGAGA | GACACTCTTCACTTGGGCCA |
| Tig1-3'UTR | CTCCCTCAAACAATCCGGCT | GTTGTCGAGCGCTTCCATTC |
| ef1-$\alpha$ | AACATCGTGGTCATCGGCCAT | GGAGGTGCCAGTGATCATGTT |
| Tig1 N.vir | AAAGTTGGAACGTGCTCTGG | CATTTGCATTGATTGGCTTG |
| ef1-$\alpha$ N.vir | TAGAGTGCAGGTGACGATCC | AGTCACCAAGTCTGCCATCA |

Supplementary Table 1
